## Supplementary material for "Functional characterisation of neuropeptides that act as ligands for both calcitonin-type and pigment-dispersing factor-type receptors in a deuterostome": Figure 1-figure supplement 1

**A**

**AjCTP1**

```
MKASLAVPITLCMFCYLLVTVTSVTINRPNAGLELSRQYPELYEKILRQLINEPTQYKRSCSNK
FAGCAHMKVANAVLKQNSRGQQQFKFGSAGPGKRSFPYLDLEEKRRVGGCGDFSGCASLKA
GRDLVRAMLRPSKFGSGGPGKK
```

**AjCTP2**

```
MKASLAVPITLCMFCYLLVTVTSVTINRPNAGLELSRQYPELYEKILRQLINEPTQEKRRVGGC
GDFSGCASLKAGRDLVRAMLRPSKFGSGGPGKK
```

**B**

**AjCTP1**

```
agctcgagctgtgtcgataacattacgcatgggaatgctatttgtcttagagtagaggga
aaaaatctcgagtcaagtcgtggagtagagttttggaaggcaatattttcaggaaaactg
tgtctgactgccgtaaaaggaacgtttttttaatttccaacaaaagagggggaagtctaa
aagccaaaaaaagagtatcttacccttgggaacaatcgttataacaaagacaacggaaggg
tcaaatcacgaaaaaccagttacagtaccattgttagcgagctaaggaacgctctttaaa
ATGAAGGCCAGTTTGGCGGTACCCATTACACTATGTATGTTTTGTTATCTACTTGTTACA
 M  K  A  S  L  A  V  P  I  T  L  C  M  F  C  Y  L  L  V  T
GTAACATCAGTAACTATTAACAGACCGAACGCCGGCTTGGAATTATCAAGACAGTATCCC
 V  T  S  V  T  I  N  R  P  N  A  G  L  E  L  S  R  Q  Y  P
GAATTATATGAAAAAATTCTTCGCCAACTTATTAACGAACCAACGCAATACAAACGGAGT
 E  L  Y  E  K  I  L  R  Q  L  I  N  E  P  T  Q  Y  K  R  S
TGCTCAAATAAGTTTGCTGGTTGTGCACACATGAAAGTCGCTAATGCAGTATTAAAACAG
 C  S  N  K  F  A  G  C  A  H  M  K  V  A  N  A  V  L  K  Q
AATAGCAGGGGTCAACAGCAATTTAAATTTGGATCGGCTGGTCCCGGAAAACGAAGTTTT
 N  S  R  G  Q  Q  Q  F  K  F  G  S  A  G  P  G  K  R  S  F
CCTTATCTTGACCTCGAAGAAAAAAGAAGGGTGGGAGGATGCGGCGATTTCAGTGGCTGT
 P  Y  L  D  L  E  E  K  R  R  V  G  G  C  G  D  F  S  G  C
GCGTCTCTTAAAGCAGGGAGAGACTTAGTTCGTGCTATGCTCCGACCGTCCAAGTTTGGG
 A  S  L  K  A  G  R  D  L  V  R  A  M  L  R  P  S  K  F  G
TCTGGGGGACCCGGCAAGAAATAGgttttattaatgaagtgtaactatagggaatctttg
 S  G  G  P  G  K  K  -
ccgcaggattaaacatttgcggctacggtcaccaccgacatacagaagacatcgtactat
gataggcctacaaaataacgttcgggaaacatcggaatggcgtatagttcctttttttgc
ttcacatgagtggatggaagtgttcaaatcatcttacagcgagtgcctaaccctccgtag
tgactgcgtcattgatgacgtcatttagccgccattagttatttcttagagctcgtttcg
ggagtaaacagaagtcttctatgacttattatgttgacaagcatcctacaattattttag
ctacggacatatctttgaaaacataaatataatatggcaattggattttgggatcccttg
taccagtttcaatcattttattccactttgtcgacttttgaatattttttttttctttt
aaattactattttaaaccaatgaagttttaggaatgctatatcataacacaaaatgtgac
gacgtcatatcttatttttttcttgtcccgaaggaaatgtttagttttcgcttccagttg
tggttttcttccttattttttgctgataatctgcaaaaaaaaaaattagttctattttaa
aactgaaacggcgtacactgaataatagtcttgaaccgaagctcgtatatgagacaggac
taatacatttcagtactatgcgtggtaaaattgaagttcctttatatatataattcaaat
atggaaggggtgggggataggaataattcaaaatcgctaataggacaatcactgcacaat
tgataacgttctattgttacaaccagttactttgaatgaaaggtcagtggaccagttttc
agggatgagcctggggtaaaaa
```

**AjCTP2**

```
agctcgagctgtgtcgataacattacgcatgggaatgctatttgtcttagagtagaggga
aaaaatctcgagtcaagtcgtggagtagagttttggaaggcaatattttcaggaaaactg
tgtctgactgccgtaaaaggaacgtttttttaatttccaacaaaagagggggaagtctaa
aagccaaaaaaagagtatcttacccttgggaacaatcgttataacaaagacaacggaaggg
tcaaatcacgaaaaaccagttacagtaccattgttagcgagctaaggaacgctctttaaa
ATGAAGGCCAGTTTGGCGGTACCCATTACACTATGTATGTTTTGTTATCTACTTGTTACA
 M  K  A  S  L  A  V  P  I  T  L  C  M  F  C  Y  L  L  V  T
GTAACATCAGTAACTATTAACAGACCGAACGCCGGCTTGGAATTATCAAGACAGTATCCC
 V  T  S  V  T  I  N  R  P  N  A  G  L  E  L  S  R  Q  Y  P
GAATTATATGAAAAAATTCTTCGCCAACTTATTAACGAACCAACGCAAGAAAAAAGAAGG
 E  L  Y  E  K  I  L  R  Q  L  I  N  E  P  T  Q  E  K  R  R
GTGGGAGGATGCGGCGATTTCAGTGGCTGTGCGTCTCTTAAAGCAGGGAGAGACTTAGTT
 V  G  G  C  G  D  F  S  G  C  A  S  L  K  A  G  R  D  L  V
CGTGCTATGCTCCGACCGTCCAAGTTTGGGTCTGGGGGACCCGGCAAGAAATAGgtttta
 R  A  M  L  R  P  S  K  F  G  S  G  G  P  G  K  K  -
ttaatgaagtgtaactatagggaatctttgccgcaggattaaacatttgcggctacggtc
accaccgacatacagaagacatcgtactatgataggcctacaaaataacgttcgggaaac
atcggaatggcgtatagttcctttttttgcttcacatgagtggatggaagtgttcaaatc
atcttacagcgagtgcctaaccctccgtagtgactgcgtcattgatgacgtcatttagcc
gccattagttatttcttagagctcgtttcgggagtaaacagaagtcttctatgacttatt
atgttgacaagcatcctacaattattttagctacggacatatctttgaaaacataaatat
aatatggcaattggattttgggatcccttgtaccagtttcaatcattttattcactttgt
cgactttttgaatattttttttttcttttaaattactattttaaaccaatgaagttttag
gaatgctatatcataacacaaaatgtgacgacgtcatatcttatttttttcttgtcccga
aggaaatgtttagttttcgcttccagttgtggttttcttccttattttttgctgataatc
tgcaaaaaaaaaaattagttctattttaaaactgaaacggcgtacactgaataatagtct
tgaaccgaagctcgtatatgagacaggactaatacatttcagtactatgcgtggtaaaat
tgaagttcctttatatatataattcaaatatggaaggggtgggggataggaataattcaa
aatcgctaataggacaatcactgcacaattgataacgttctattgttacaaccagttact
ttgaatgaaaggtcagtggaccagttttcagggatgagcctggggtaaaaa
```
