## Supplementary material for "Functional characterisation of neuropeptides that act as ligands for both calcitonin-type and pigment-dispersing factor-type receptors in a deuterostome": Figure 1-figure supplement 2

```
A_japCT1   -----SCSNKFAG-CAHMKVANAVLKQNSRGQQQFKF--GSAG--Pa
A_japCT2   --RVGGCGD-FSG-CASLKAGRDLVRAMLR---PSKF--GSGG--Pa
H_scaCT1   -----SCSDRFSG-CAHLKVAKALLDQ-ARREENSRF--GISG--Pa
H_scaCT2   ---MGGCGD-FSG-CASLKAGRDLVRAMLRQ--PSKF--GSGG--Pa
H_glaCT1   -----SCSDRFAG-CTHLKLAKVLIDQ-SRRDGNIRF--GSSG--Pa
H_glaCT2   ---MGGCGDKFSG-CASLKAGRDLVRAMLRQ--PSKF--GSGG--Pa
S_purCT    ---SKGCGS-FSG-CMQMEVAKNRVAALLRNSNAHLF--GLNG--Pa
A_rubCT    NGESRGCSG-FGG-CGVLTIGHNAAMRMLAES-NSPF--GASG--Pa
A_plaCT    --SSSGCAEYFGG-CAQLKLGQDALSRMLADS-NSRF--GSGG--Pa
P_pecCT    --SGTGCTQ-FSG-CAQLKVGQDALSRVLADS-NSRF--GSGG--Pa
O_vicCT1   -SGNGGCAG-FTG-CAQLAAGQNALRNFMHSNRASLFT-GASG--Pa
O_vicCT2   -NGNGGCAG-FTG-CAQLAAGQSALQAMIHSGRASLF--GSGG--Pa
O_araCT1   -GTEKGCSG-FSG-CAQLAAGQSALQAMIHGNRASLF--GSGG--Pa
O_araCT2   --GGGGCKS-FSG-CAQLVIGQNAVRNMMHSNRASIFS-GPRG--Pa
A_filCT1   -GTEKGCAG-FSG-CAQLAAGQSALQAMIHGNRASLF--GSGG--Pa
A_filCT2   ---GGGCRG-FSA-CAQLAIGQDAFRNMMHNNRASIFT-SARG--Pa
A_medCT    ----G-C-DVFGG-CAQLKVGRELAIDTLQKGSGGMF--GSSG--Pa
An_japCT   ----GDC-DSFGG-CAQLTASQQLAYRTRHRDLSNLF--GTGG--Pa
S_kowCT1   ---GEGCKG-FAA-CGNLDAGRKHWKANMGGGRTAMESSASGT--Pa
S_kowCT2   GRGTSACGG-FAT-CKQLEYGRKYATS---KADLSHF--GATS--Pa
B_floCT1   --GKIACKT--AW-CMNNRLSHNLSSLDNPTDTGVG------A--Pa
B_floCT2   -----DCST--LT-CFNQKLAHELAMDNQRTDTANPY-----S--Pa
B_floCT3   -----KCES--GT-CVQMHLADRLRLGLGHNMFTNT---GPES--Pa
C_intCT    ------CDG-VST-CWLHELGNSVHATAGGKQ-NVGF-----G--Pa
H_sapCGRP  -----ACDT--AT-CVTHRLAGLLSRSGGVVKNNFVPT-NVGSKAFa
H_sapCT    ------CGN-LST-CMLGTYTQDFNKFHTFP--QTAI--GVGA--Pa
L_gigCT    -----SCSLRLGGMCLTENLNAAANQY-------EYLS-SGLS--Pa
D_melDH31  -TVDFGLARGYSG---TQEAKHRMGLA------AANF---AGG--Pa
C_telDH31  -----RFDAGYGS---RYGVAQSVGSKLMALKQAADW----NG--Pa
C_telCT    -----TCQFNLGGHCATESAASVADHW-------HYLN-SPLS--Pa
```
