## Supplementary figures and images for "Functional characterisation of neuropeptides that act as ligands for both calcitonin-type and pigment-dispersing factor-type receptors in a deuterostome"

### Figure 2-figure supplement 1

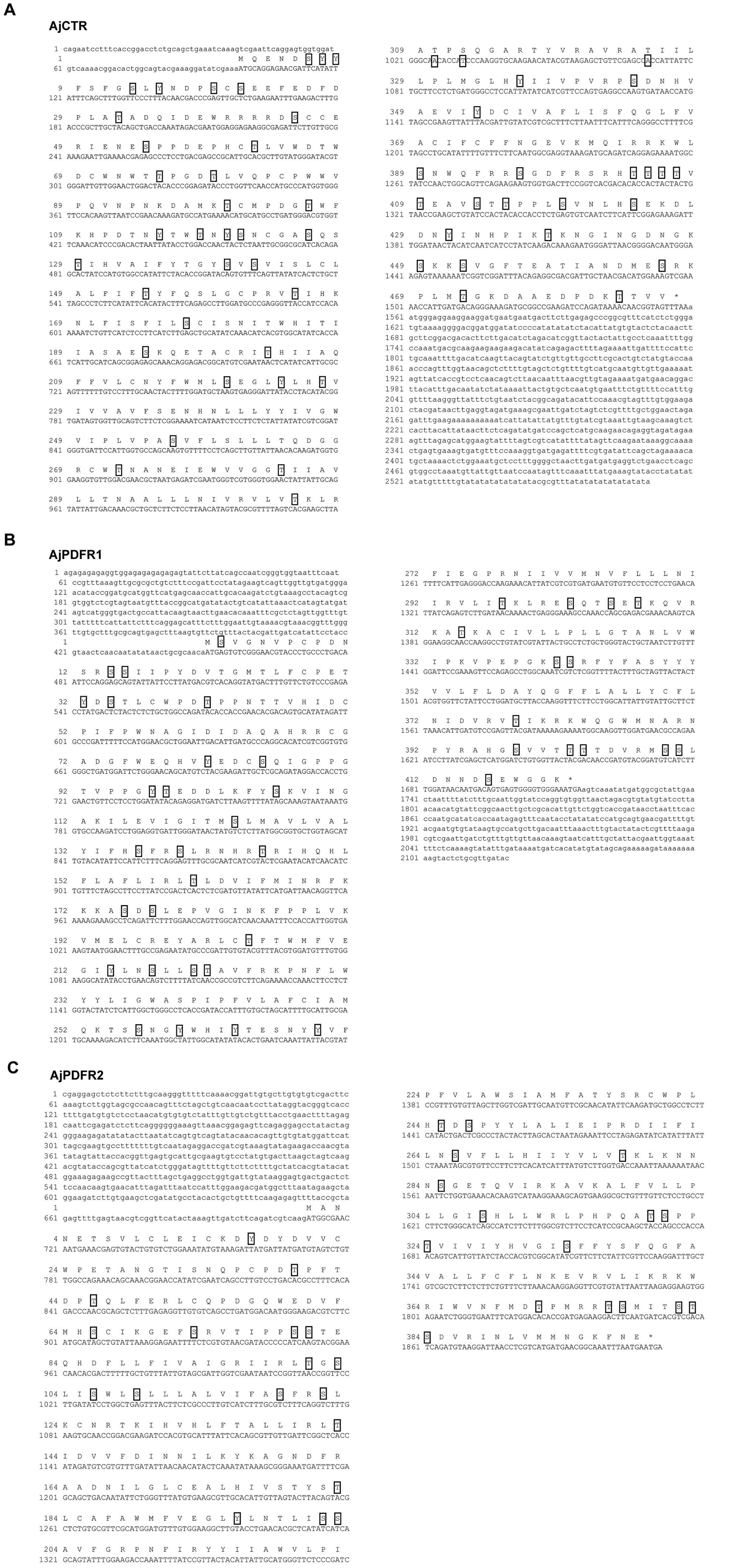

### Figure 2-figure supplement 2

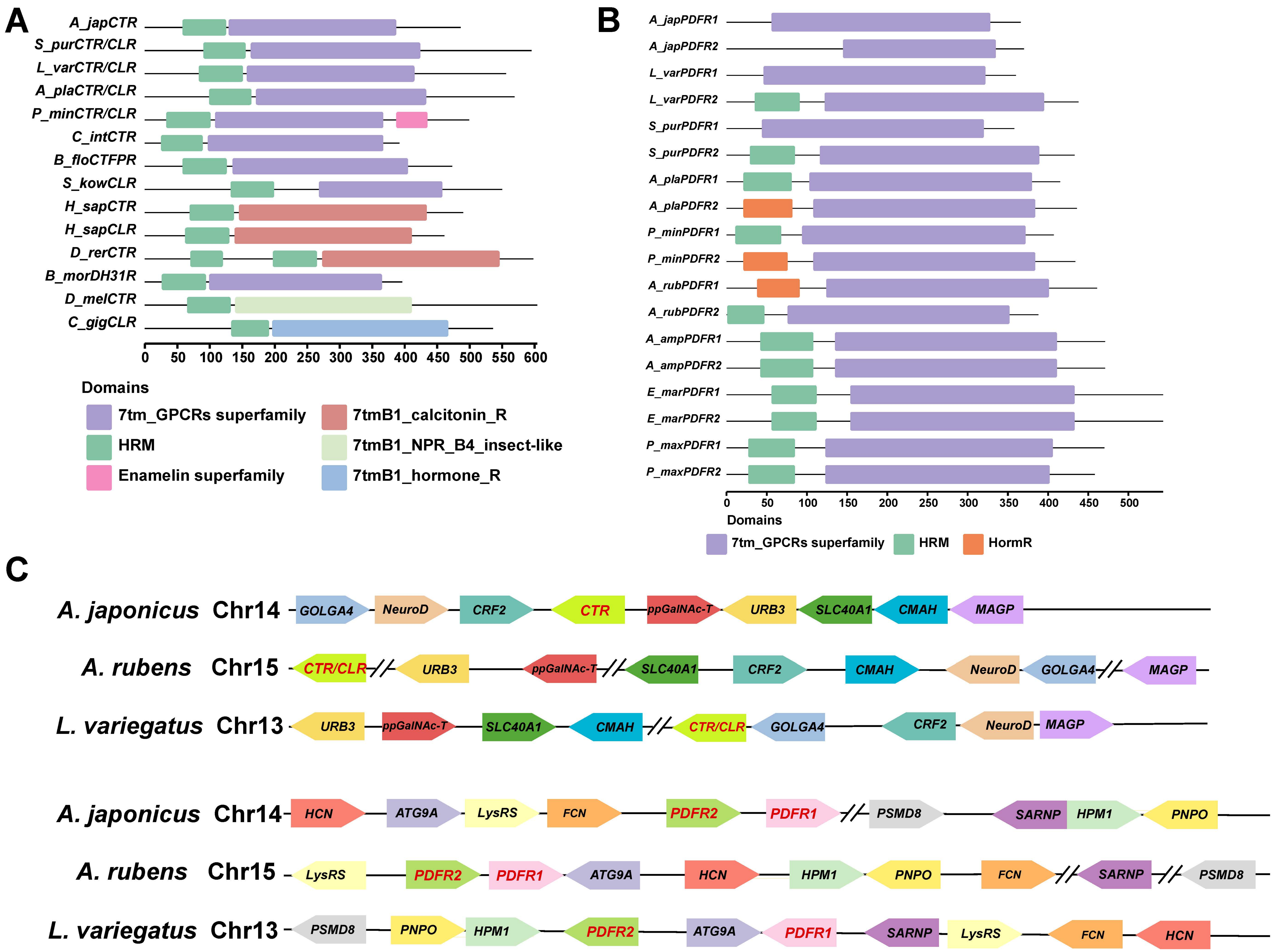

### Figure 4-figure supplement 1

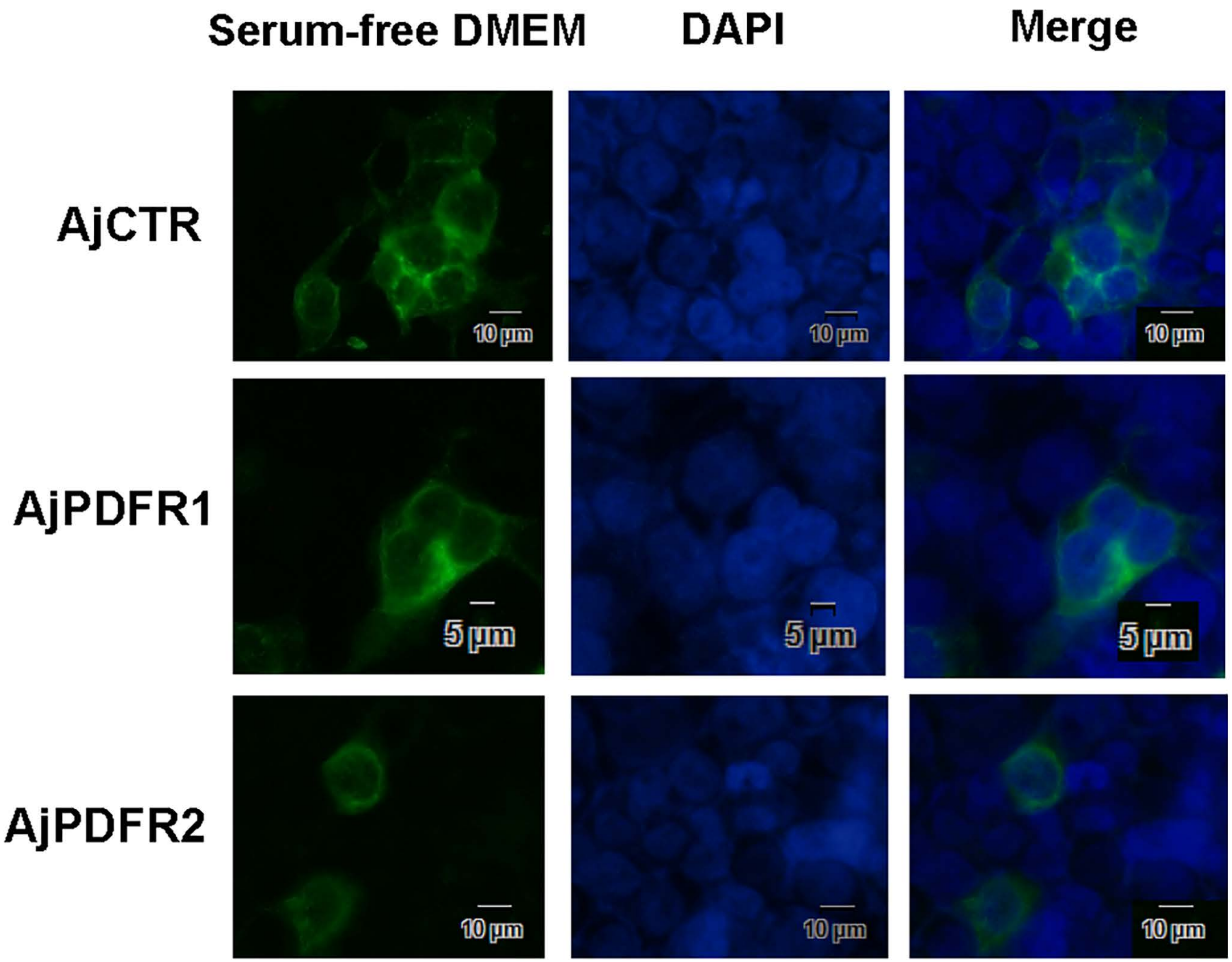

### Figure 5-figure supplement 1

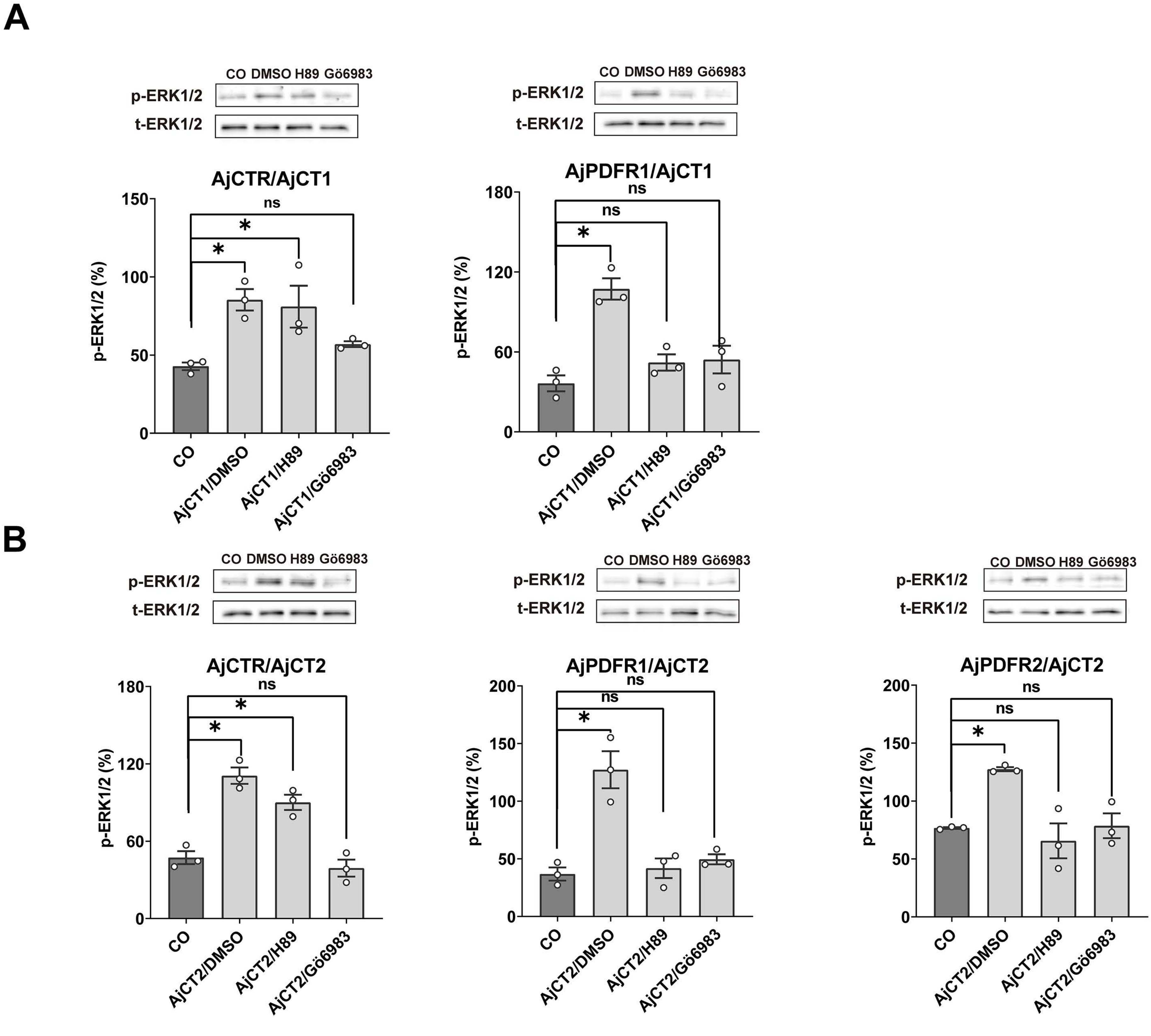

### Figure 7-figure supplement 1

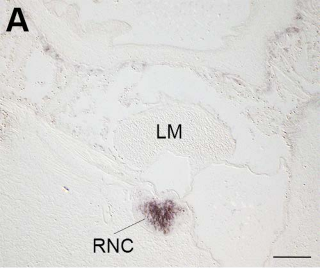

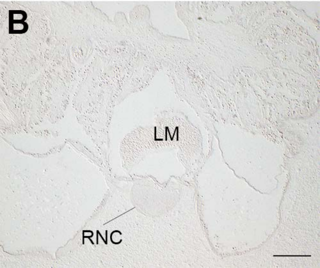

### Figure 8-figure supplement 1

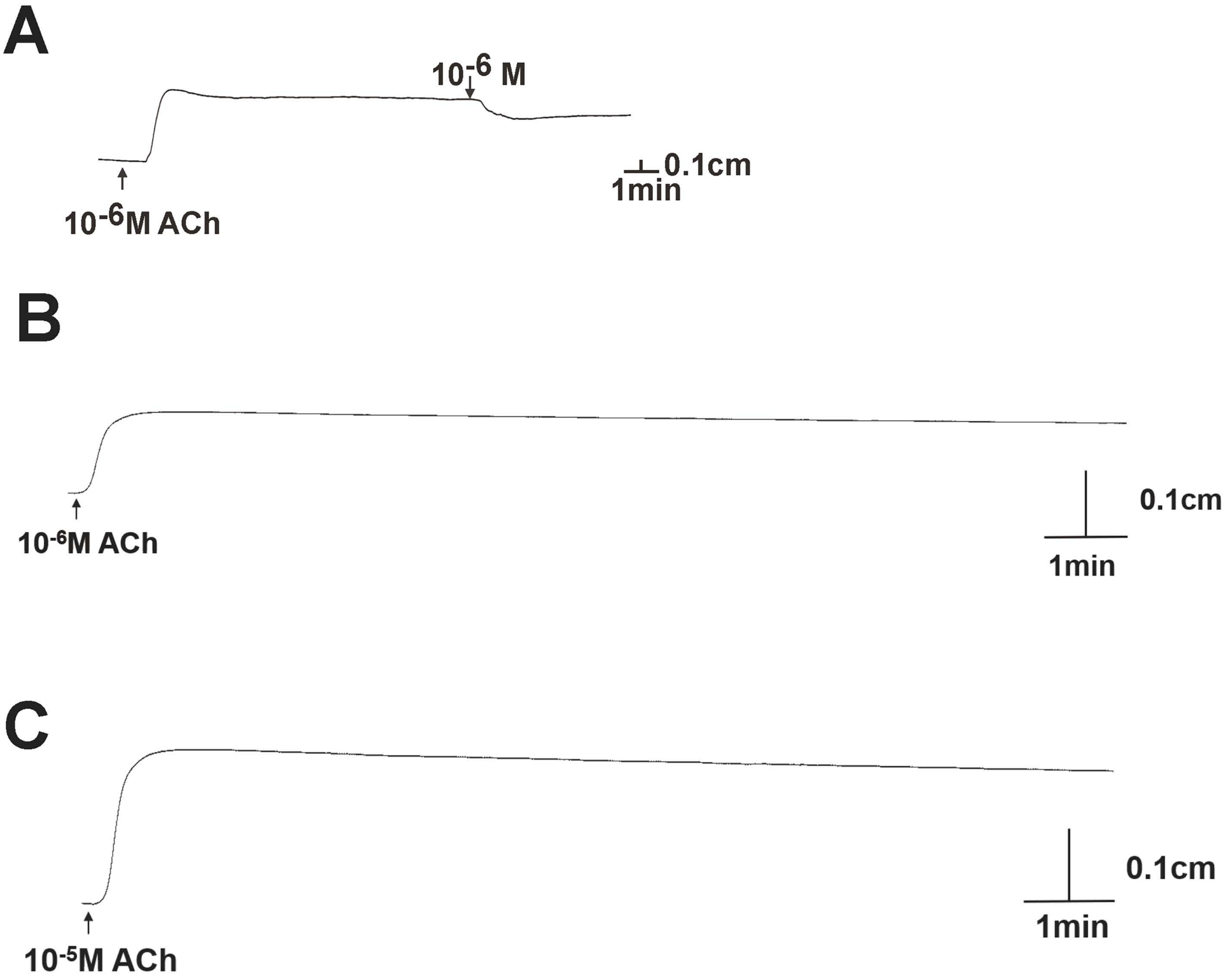

### Figure 9A-figure supplement 1

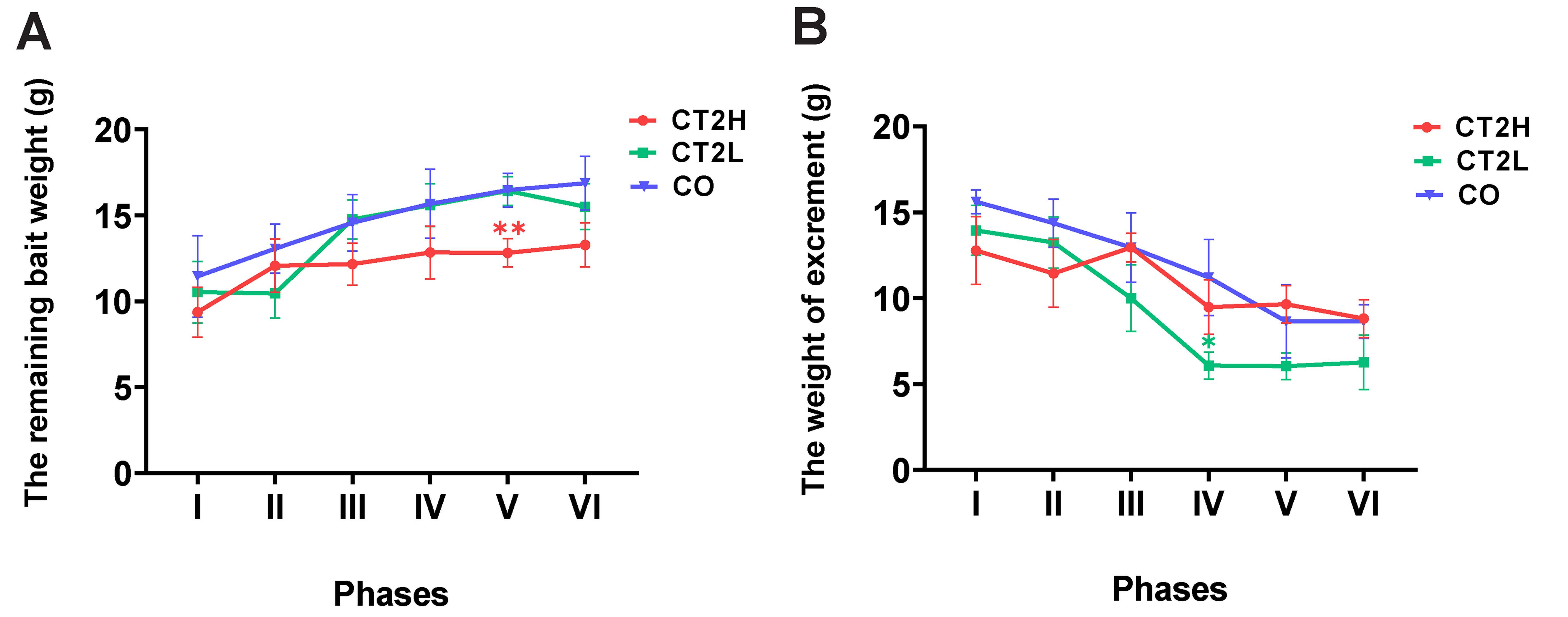

### Figure 10-figure supplement 1

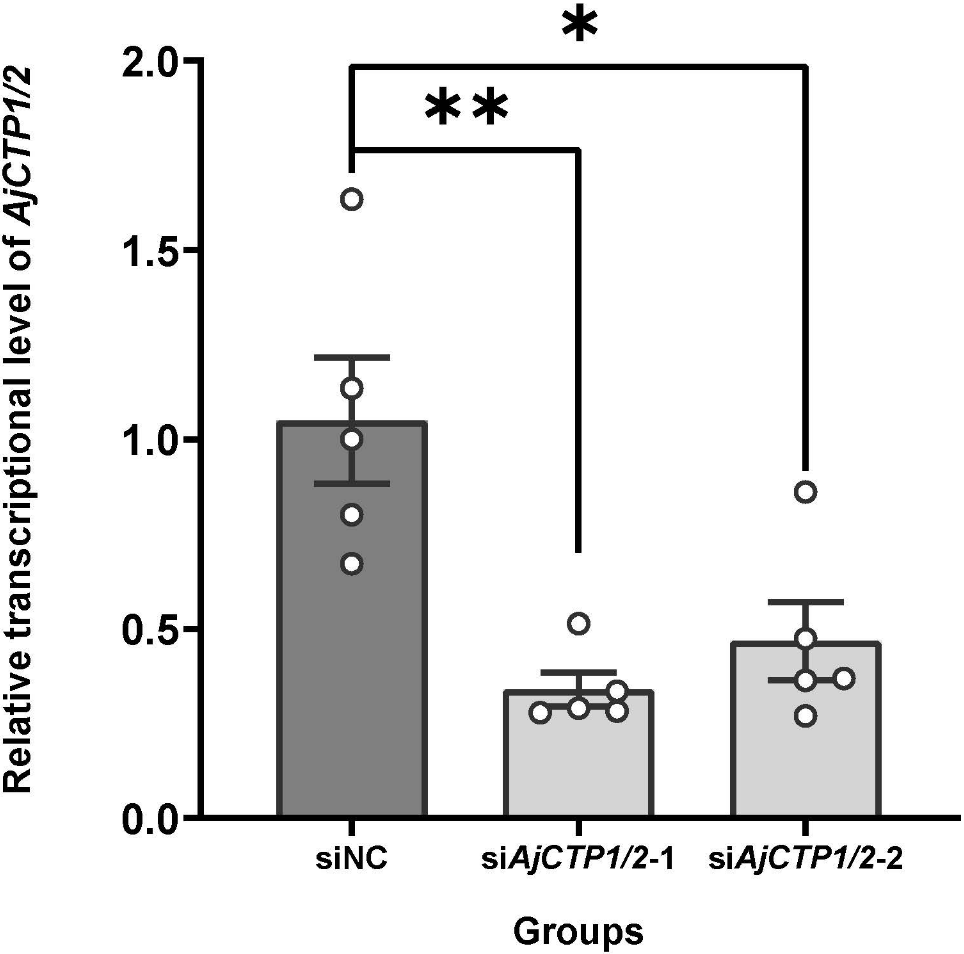
